# *In vivo* base editing rescues photoreceptors in a mouse model of retinitis pigmentosa

**DOI:** 10.1101/2022.06.20.496770

**Authors:** Jing Su, Kaiqin She, Li Song, Xiu Jin, Ruiting Li, Qinyu Zhao, Jianlu Xiao, Danian Chen, Hui Cheng, Fang Lu, Yuquan Wei, Yang Yang

**Author notes:** Corresponding authors: Fang Lu, Postal address: Guoxue xiang, No.37, Chengdu, Sichuan, China 610041, Yang Yang, Postal address: Ke-yuan Road 4, No. 1, Gao-peng Street, Chengdu, Sichuan, 610041, China. These authors contributed equally: Jing Su, Kaiqin She, Li Song.

## Abstract

Retinitis pigmentosa (RP) is a group of retinal diseases that cause the progressive death of retinal photoreceptor cells and eventually blindness. Mutations in the β-domain of the phosphodiesterase 6 (*Pde6b*) gene are among the most identified causes of autosomal recessive RP. Here, we report a base editing approach in which adeno-associated virus (AAV)-mediated adenine base editor (ABE) delivery to postmitotic photoreceptors is used to correct the *Pde6b* mutation in a retinal degeneration 10 (*rd10*) mouse model of RP. Subretinal delivery of AAV8-ABE corrects *Pde6b* mutation with up to 37.41% efficiency at the DNA level and up to 91.95% efficiency at the cDNA level, restores PDE6B expression, preserves photoreceptors and rescues visual function. RNA-seq reveals upregulation of genes associated with phototransduction and photoreceptor survival. Our data demonstrate that base editing is a potential gene therapy that could provide durable protection against RP.

## Introduction

Retinitis pigmentosa (RP) is the leading cause of progressive vision loss and inherited blindness and is characterized by degeneration of photoreceptors, affecting approximately 1 in 4000 people worldwide(*1*). Most patients with RP manifest with night blindness due to loss of rods, followed by daytime vision loss due to secondary degeneration of cones(*2*). RP is a genetically heterogeneous disease caused by mutations in more than 50 genes, including autosomal recessive, autosomal dominant, X-linked, and mitochondrial inheritance patterns(*3*). The most common form of autosomal recessive RP is associated with mutations in the β-subunit of rod cGMP-phosphodiesterase (PDEβ), which is encoded by the *Pde6b* gene(*4*). *Pde6b-*RP accounts for 4-5% of autosomal RP and is one of the earliest-onset and most severe forms of the disease(*5*).

Currently, there is no effective treatment that can stop the progression of RP and restore the vision. Clinically, the management of RP is restricted to slowing down the degeneration process by low vision aids, vitamin therapy and sunlight protection(*6*). Therefore, scientists are working hard to find a cure for RP. Several RP-related clinical trials are underway, such as stem cell therapy by jCyte, gene therapy for *Pde6b* mutation by Coave, and *Nr2e3* gene therapy trials initiated by Ocugen. The therapeutic results of these trials, however, have not yet been announced. In addition, many therapeutic strategies have been developed to rescue retinal photoreceptor degeneration in preclinical animal models of RP, such as the introduction of small molecules for promoting cell survival(*7–9*), genetic complementation *via* viral-vector injection(*10, 11*), gene editing by electroporation *in vivo*(*12, 13*), or stem cell transplantation(*14*). However, each of these approaches shows drawbacks, including limited efficiency or lack of durable gene correction to rescue phenotypes.

In recent years, a base editing technique has been developed to convert individual base with exposed non-target strands through the action of cytosine or adenosine deaminase coupled to the CRISPR-associated protein 9 (Cas9) complex(*15*). Base editing can directly convert targeted base pairs without generating DNA double-strand breaks (DSBs) and with minimal indels, so it is considered more suitable for the treatment of human monogenetic disorders(*16*). The adenine base editors (ABEs), which is composed of dCas9 and adenine deaminase, can convert A•T to G•C or C•G to T•A. In recent years, studies have reported that lentiviral vector-delivered ABE could correct *Rpe65* mutation in the retinal pigment epithelium of *rd12* mice, a mouse model of Leber congenital amaurosis (LCA)(*17, 18*). To explore whether base editing can effectively rescue the dysfunction of photoreceptors and provide a potential therapeutic strategy for RP, we aimed to correct the *Pde6b* mutation in the *rd10* mouse model, which recapitulates the human condition and is analogous to the mutations that cause human RP(*19, 20*).

In this study, we developed a strategy using ABE based on AAV8, which has high retina tropism, to correct the pathogenic point mutation in *rd10* mice. We found that subretinal delivery of AAV8-ABE corrected the *Pde6b* mutation in the photoreceptors, preserved rods and cones and restored visual function in *rd10* mice, suggesting a therapeutic opportunity for base editing in the treatment of patients with RP.

## Results

### *In vitro* screening of ABEs targeting *Pde6b*-R560C

Similar to a human RP, *rd10* mice are recessively inherited and carry a missense mutation (c.1678C to T, R560C) in the *Pde6b* gene (Fig. 1A). To evaluate the base editing efficiency *in vitro*, we generated a HEK293-*Pde6b* mutant cell line by stably integrating the *rd10* mutant *Pde6b* sequence into the AAVS1 genomic locus using CRISPR/Cas9(Fig. S1 and Table S1). We first searched for protospacer sequences that span the targeted base. Since the targeting site of the *rd10* mice lacks the canonical NGG protospacer-adjacent motif (PAM) sequence and ABEs favorably deaminate within the window of protospacer positions 4-8(*21*), we designed two sgRNAs (sgRNA-A6, sgRNA-A8) with the NG PAM sequence suitable for most ABEs (Fig. 1A). XCas9(3.7)-ABE (x7.10), xABEmax, NG-ABEmax, NG-ABE8e and SpG-ABE and modified SpG-ABE8e are compatible with the PAM sequences of sgRNA-A6 and sgRNA-A8. We co-transfected various combinations of ABE and sgRNA into HEK293-*Pde6b* cells to screen the optimal ABE and sgRNA and performed sequencing analysis 72 hours after transfection. Sanger sequencing showed that the on-target editing efficiency of NG-ABE8e co-transfected with sgRNA-A6 and sgRNA-A8 was higher than that of other combinations (31.33±4.73% and 34.33±2.08%, respectively) (Fig. 1B). However, both NG-ABE8e and SpG-ABE8e co-transfected with sgRN-A6 showed a higher rate of bystander A9 to G conversion than sgRNA-A8 (25.33±3.79% and 22.67±1.53%, respectively) (Fig. 1B). The highest on-target editing efficiency was observed with NG-ABE8e co-transfected with sgRNA-A8, which was then selected for further studies (Fig. 1B, 1C).

**Fig. 1.**
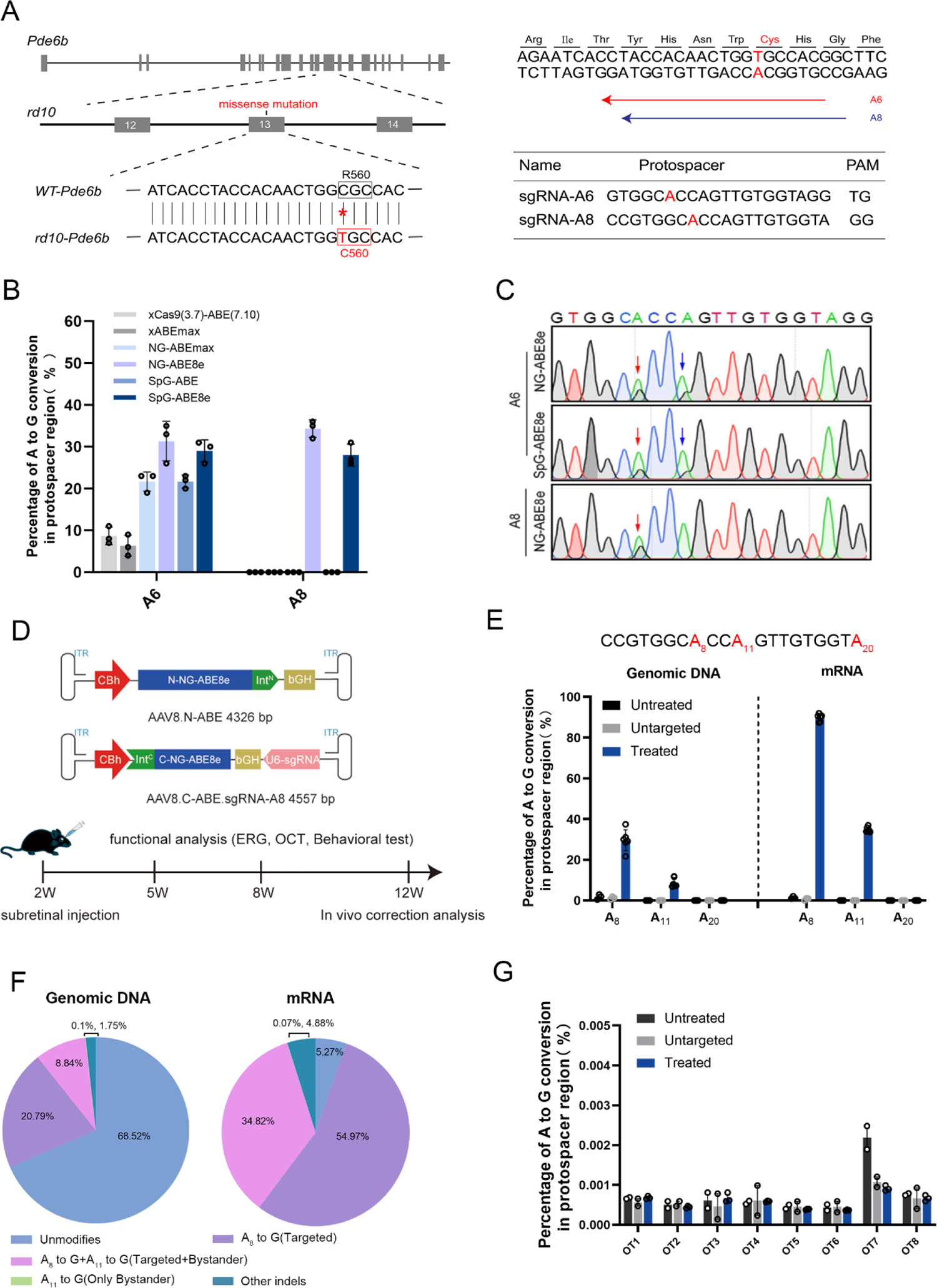
*In vitro* screening of optimal ABE and sgRNA and *in vivo* validation of *Pde6b* mutation correction by AAV8-ABE. (**A**) Schematic representation of a missense mutation in the *rd10* mouse *Pde6b* gene (c.1678C to T, R560C) that causes RP (left). sgRNA-A6 and sgRNA-A8 were designed to target the mutation site (red letter) in the editing window of ABEs (right). (**B**) Sanger sequencing quantifies the correction efficiency of ABEs in mutant cell lines (n=3 biological replicates each). Mean ± SD are shown. ( **C**) Representative direct genome sequencing results of gene correction from NG-ABE8e/SpG-ABE8e cotransfection with sgRNA-A6 or sgRNA-A8. Red arrows indicate correction of the target site and blue arrows indicate bystander editing. (**D**) Schematic diagram of the genomes of two AAV viral vectors encoding split-intein NG-ABE8e (top) and a summary of the *in vivo* experiments (bottom). (**E**) Correction efficiency at the DNA and cDNA levels in *rd10* mice after base editing treatment. Untreated and untargeted *rd10* mice (n=3, each group) were included as controls. Treated *rd10* mice (n=6). Mean ± SD are shown. ( **F**) The pie charts show the average composition of allele variants at the DNA (left) and cDNA levels (right) in treated *rd10* mice. (**G**) NGS analysis of the top eight potential off-target sites in retinal DNA samples. Untreated and untargeted *rd10* mice (n=2, each group), treated mice (n=3). Mean ± SEM are shown.

### *In vivo* base editing corrects the *Pde6b* mutation in the *rd10* mice

To efficiently deliver ABE and sgRNA to mouse retinal tissue *in vivo,* AAV8 was used to deliver the base editor system. Considering the limitation of AAV package capacity, we engineered a split-intein dual-AAV8 system to express the amino segment of ABE and the carboxyl terminus connected to sgRNA/untargeted control sgRNA in two distinct vectors (refer to as AAV8.N-ABE, AAV8.C-ABE.sgRNA-A8 and AAV8.C-ABE.sgRNA-ctrl, respectively) (Fig. 1D). We performed subretinal injection with AAV8 vectors in *rd10* mice at 2 weeks of age (AAV8.N-ABE and AAV8.C-ABE.sgRNA-A8 as a treated group, AAV8.N-ABE and AAV8.C-ABE.sgRNA-ctrl as a control untargeted group, 3×10 ^9^ GC per eye for each AAV8). Since AAV-mediated gene expression takes time, the *rd10* mice were raised in darkness until 4 weeks old to extend the treatment window(*22*), and the therapeutic effect in mice was assessed at several time points (Fig. 1D).

Genomic DNA and mRNA were extracted from mouse retinal tissue to quantify the correction efficiency at 12 weeks of age (10 weeks post-injection). Next-generation sequencing (NGS) results showed that treated retinas had an average correction rate of up to 29.63% at the DNA level (range 21.67% ∼ 37.41%, n=6 eyes). In addition to targeted A6 editing, we also detected approximately 8.84% of bystander editing occurred at A11 site (Fig. 1E). As in our previous reported study(*23*), we observed higher correction efficiency at the cDNA level, averaging up to 89.79% (range 87.41% ∼ 91.95%, n=6 eyes), accompanied by the production of 34.82% bystander editing (Fig. 1E). When we examined the percentage of precise correction edits without bystander editing, the correction efficiencies at the DNA and cDNA levels were still as high as 20.79% and 54.97%, respectively, while bystander editing alone were both less than 0.5% (Fig. 1F). We also further performed NGS analysis on the top eight potential off-target sites for sgRNA-A8 identified by the algorithm described in www.benchling.com. Samples from treated animals did not show indel rates above background, suggesting that sgRNA-A8 specifically targets the intended sites (Fig. 1G).

In addition, we found that further increasing the injection dose (a total of 6×10 ^10^ GC/eye) did not improve the correction rate of the target site (average 31.30%), but with higher bystander production at the A11 site (average 15.00%) (Fig. S2). We infer that the functional base editor acting on the target site had reached saturation at the dose of 6×10 ^9^ GC/ eye, making high-dose injection unnecessary.

### *In vivo* base editing restores PDE6B expression and prolongs photoreceptor survival

To explore whether the corrected *Pde6b* gene could be translated into PDE6B protein, we performed Western blot (WB) on the retinas of wild-type (WT) mice, and untreated, untargeted and treated *rd10* mice. The results showed that PDE6B protein was detectable in the retinas of the treated mice, while it was completely absent in the untreated and untargeted mice (Fig. S3). Meanwhile, the rod outer segment (OS) of treated retinas exhibited PDE6B immunolabeling, demonstrating that base editing restored PDE6B expression (Fig. 2A). To analyze the effect of base editing on photoreceptor survival, the retinal sections were immunolabeled with rod OS and cone specific antibodies. The retinas of treated mice showed remarkable preservation of rods and cones, whereas the photoreceptors completely degenerated in untreated and untargeted mice (Fig. 2A). To quantify rods and cones rescue, we measured rod OS length and cone numbers. As expected, the rod OS length of treated *rd10* mice was significantly shorter than that of WT mice (6.3±2.5 µm vs 19.3±1.5 µm), because the rod degeneration began prior to treatment (Fig. 2B). However, the number of arrestin-stained cones was significantly increased compared with that in the untreated group, corresponding to about 89% of WT mice (11.4±1.5 vs 12.8±1.3 per 100 µm) (Fig. 2C). To further determine M-cone and S-cone rescue, we performed immunostaining of retina flatmounts. The results revealed that M-cones and S-cones in the treated group were almost comparable to those in WT mice, showing remarkable photoreceptor preservation compared to untreated mice (Fig. S4).

**Fig. 2.**
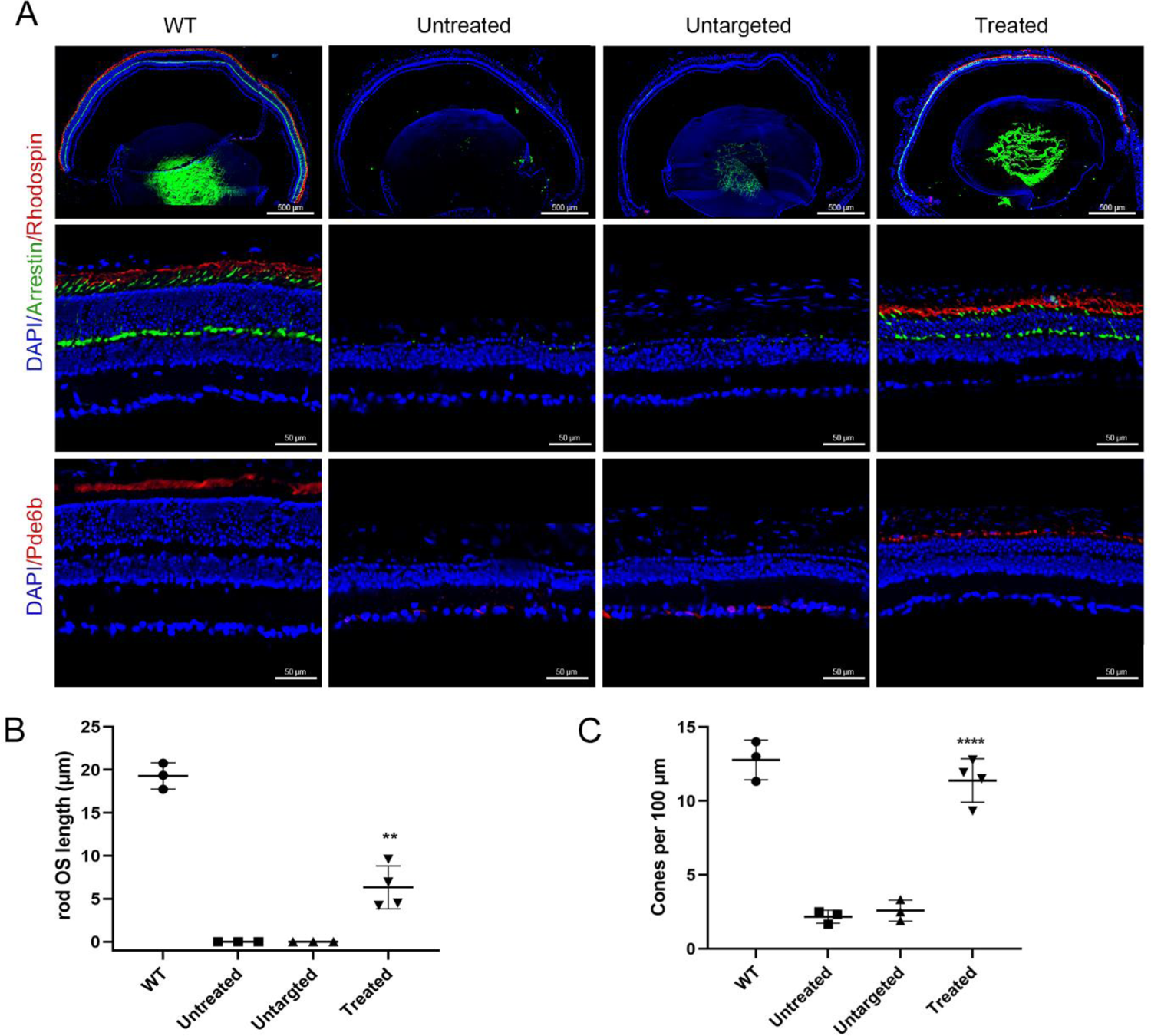
*In vivo* base editing restores PDE6B expression and prevents photoreceptor degradation. (**A**) Top and middle: representative retinal cross-sections of 12-week-old mice from each group were immunolabeled for rhodopsin (Rho, rod cells, red) and arrestin (cone cells, green). Nuclei are labeled with DAPI (blue). Scale bars: 500 μm for low power; 50 μm for high power. Bottom: Representative PDE6B immunostaining (red) of retinal sections from 12-week-old mice in each group. Scale bars: 50 μm. (**B, C**) Quantitative results of rod OS length (B) and cone number (C) from WT mice(n=3), untreated mice(n=3), untargeted mice (n=3) and treated mice (n=4). Mean ± SD are shown. Comparison between treated and untreated groups, **p<0.01, ****p<0.0001, one-way ANOVA analysis with Tukey’s post-hoc test.

### *In vivo* base editing rescues retina and RPE structure

Degeneration of photoreceptors in *rd10* mice results in structural changes in the retina. We visualized and assessed the structure of retina *in vivo* using optical coherence tomography (OCT). The retinal thickness of treated mice was significantly preserved compared to untreated mice, as seen by OCT taken at 5 weeks of age and continuing to 12 weeks of age (Fig. S5A). At 12 weeks of age, the outer nuclear layer (ONL) and inner segment (IS)/OS layer of the untreated or untargeted eyes were barely visible in OCT (Fig. 3A), which was confirmed by transmission electron microscopy (TEM) (Fig. 3C). In contrast, the ONL and IS/OS layer partially remained in treated eyes (Fig. 3A, 3C). The thicknesses of the whole retina and ONL were measured in the OCT scan across the optic disc at multiple positions, and quantitative analysis showed that the retinal and ONL thicknesses were 139.0∼153.4 µm and 20.6∼26.8 µm in treated mice, respectively, which were significantly thicker than those in untreated mice, equivalent to approximately 62.6%∼66.7% and 35.8%∼47% of WT, respectively (Fig. 3B). Hematoxylin and eosin (H&E) staining also showed that the ONL in untreated *rd10* mice was reduced to only one layer of incomplete nuclei, whereas in treated mice the ONL was rescued and maintained at 4-6 layers of nuclei in more than 3/4 area of the retina, as with the above immunohistochemical staining (Fig.S6).

**Fig. 3.**
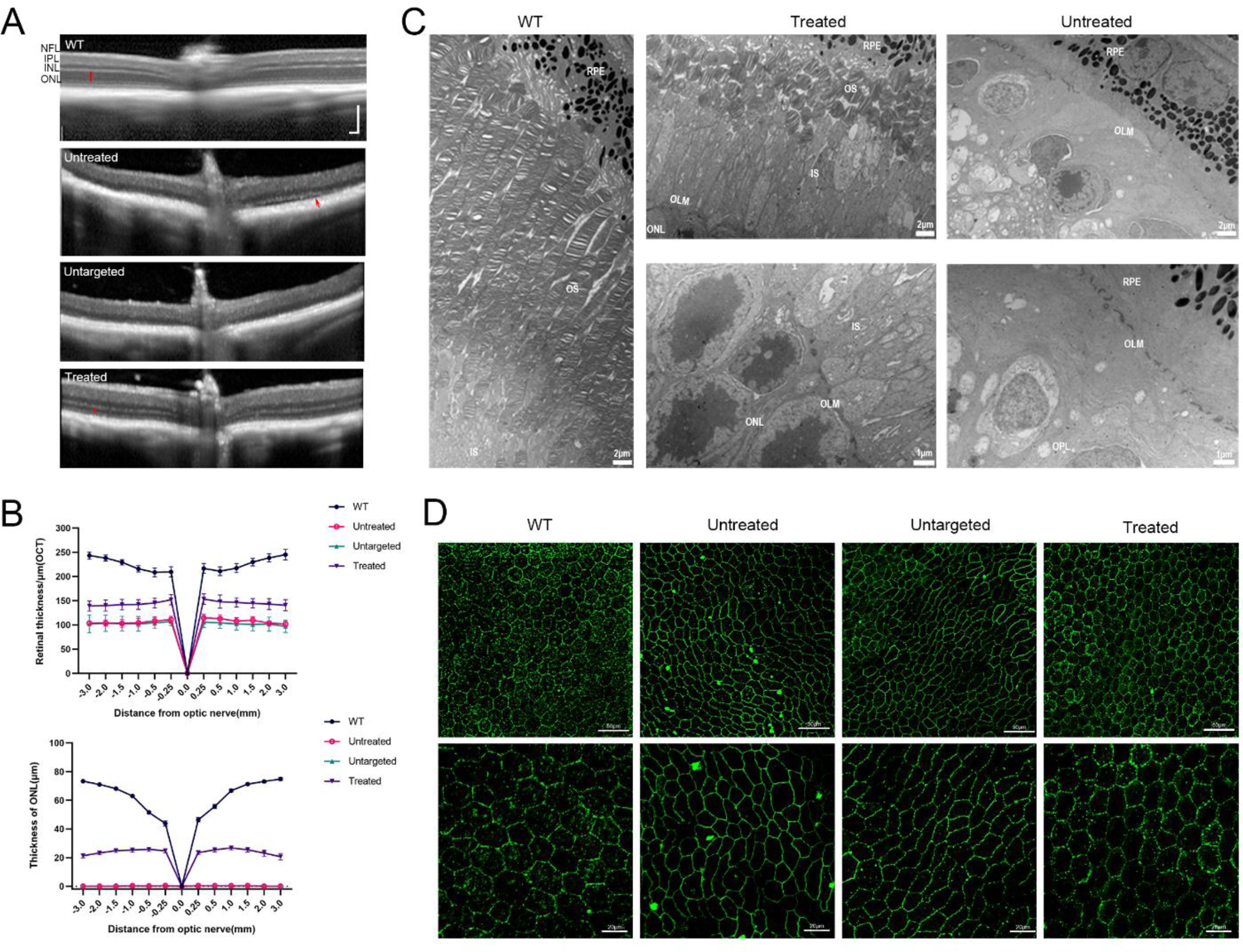
Retinal and RPE structures are rescued in treated *rd10* mice after base editing. (**A**) Representative OCT images from 12-week-old WT mice, untreated, untargeted and treated *rd10* mice. The red bar marks the ONL layer. The red arrow indicates the separation of the retina from pigment epithelial cells. (**B**) The results of retinal (top) and ONL thickness (bottom) measurements at different distances from the optic disc (n=10, each group). Mean ± SEM are shown. ( **C**) Ultrastructural analysis of retina by TEM. The OS, IS and ONL layers are clearly visible in both WT and treated mice, whereas untreated mice have degenerated. Scale bars: 2 μm for low power; 1 μm for high power. (**D**) Representative images of flat-mounted RPE with ZO-1 immunostaining from each group. The morphology of untreated/untargeted RPE cells became distorted, elongated and lost the original regular hexagon. Scale bars: top, 50 μm; bottom, 20 μm.

Collateral damage from photoreceptor degeneration in *rd10* mice causes changes in retinal pigmented epithelium (RPE) cell morphology(*24, 25*). Flat-mounted RPE was stained with ZO-1 antibody to evaluate the rescue of RPE in treated mice. We observed that the morphology of untreated RPE cells became distorted, elongated and lost the original regular hexagon compared to WT mice at 12 weeks of age. However, regular hexagon shapes were detected in the treated group, indicating that base editing can rescue the structure of the RPE in *rd10* mice (Fig. 3D).

### *In vivo* base editing restores retinal function in the *rd10* mice

Severe and progressive photoreceptor degeneration in *rd10* mice results in a complete absence of electroretinography (ERG) responses at 2 months after birth(*26*). To determine whether base editing can restore retinal function, we performed scotopic (rod-driven) and photopic (cone-driven) ERG recordings in mice at 12 weeks of age. As shown in Fig. 4A, we found that the treated *rd10* mice had substantial retinal responses, with both a- and b-waves of ERG clearly visible. The a- and b-waves of the ERG represent the photoreceptor response and bipolar cell activity, respectively. At the 2.2 log cd s/m^2^ of stimulus intensity under scotopic conditions, the treated *rd10* mice retained a-wave and b-wave amplitudes that were approximately 11.37% and 29.50%, respectively, of the ERG amplitudes from WT mice (a-wave:13.06±0.56 µV vs 114.72±9.76 µV; b-wave:79.28±10.41 µV vs 268.73±23.19 µV), whereas untreated mice and untargeted mice did not show any detectable ERG signals (Fig. 4A and Fig. S7A). Photopic ERGs were recorded at two wavelengths to assess photopic retinal vision, which is mediated by two types of cones with different wavelength sensitivities: M-cones are optimally activated under green light, while S-cones are more sensitive to UV light. Our data showed that both photopic ERG waveforms from the treated *rd10* mice were significantly improved compared to the untreated or untargeted mice (Fig. 4B, 4C). At the 3.9 log cd s/m^2^ of stimulus intensity under photopic conditions, the a- and b-waves of M-cone-dependent ERG were approximately 16.67% and 23.27% of those of WT mice (a-wave:13.72±2.65 µV vs 87.27±4.16 µV; b-wave:54.38±6.08 µV vs 233.68±9.01 µV), respectively. Meanwhile, the a- and b-waves of S-cone-dependent ERG were 11.73±2.7 6 µV and 65.37±9.80 µV, respectively, corresponding to about 13.95% and 26.65% of ERG waveforms in WT mice (84.09±5.51 µV; 245.24±12.05 µV) (Fig. 4B, 4C and Fig. S7B). From 5 to 12 weeks of age, the amplitude of ERG in the untreated group had entirely fallen to zero, whereas it maintained approximately 50% in the treatment group, indicating that the *in vivo* base editing restored partial retinal function (Fig. 4D, 4E).

**Fig. 4.**
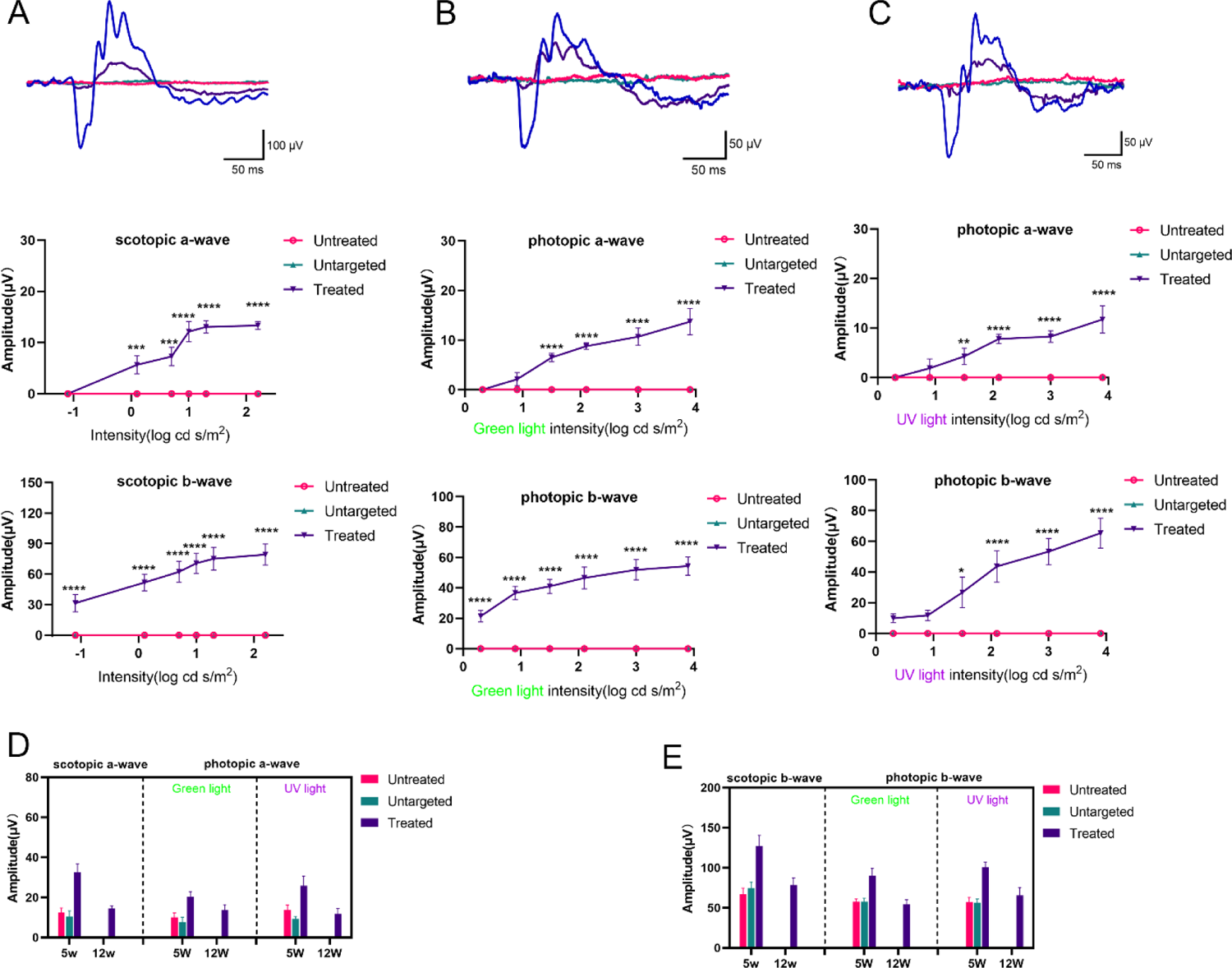
Scotopic and photopic ERG responses were enhanced in treated *rd10* mice. (**A**) Representative scotopic ERG waveforms at a light stimulus of 2.2 log cd s/m^2^ (top) and quantification of scotopic ERG a- and b-waves at 12 weeks of age in mice (middle and bottom). Blue, WT mice; red, untreated *rd10* mice; green, untargeted *rd10* mice; purple, treated *rd10* mice. (**B, C**) Representative photopic ERG waveforms at green/UV light stimulus of 3.9 log cd s/m^2^ (top) and quantification of photopic ERG a- and b-waves (middle and bottom). (**D, E**) Quantification of scotopic and photopic ERG a- and b-waves in mice at different time points at 2.2 log cd s/m^2^ and 3.9 log cd s/m^2^ stimulus intensity, respectively. (**A-E**) Untreated and untargeted *rd10* mice (n=10, each group) were included as controls. Treated *rd10* mice (n=10). Mean ± SEM are shown. Comparison between treated and untreated groups, *p<0.05, **p<0.01, ***p<0.001, ****p<0.0001, two-way ANOVA with Tukey’s post-hoc test.

### *In vivo* base editing improves visual behavior in *rd10* mice

To test whether rescue of retinal function and structure through base editing can translate into improvements in vision, we used two modified visual cliff tests to assess visually guided behavior. The two tests relied on the animals’ innate tendency to avoid the deep side of a visual cliff field (the “deep side” is also known as the unsafe zone; the other shallow side is the safe zone). The first test determined the mice’s preference for safe or unsafe zone by placing the mice on a pedestal between safe and unsafe zone; the second test measured the time that the mice spent in the safe and unsafe zone, known as the cliff open field test (Fig. 5A, 5B). In the 10 step-down trials, the mean number of choices in the safe zone was 7.0±0.5 for the WT mice and 6.3±0.8 for the treated mice at 12 weeks of age, both showing a preference for the safe side. In contrast, the untreated mice did not prefer either side, with an average of 4.4±0.5 and 4.3±0.4 selections in the safe and unsafe zone, respectively (Fig. 5C). Corresponding to the results of the step-down trials, 12-week-old untreated mice spent comparable time in the safe zone (152.8±17.4 s) and the unsafe zone (147.2±17.4 s), and the treated mice spent more time in the safe zone (179.0±16.2 s vs 121.0±16.2 s), but there was no significant difference compared to untreated mice (Fig. 5D, 5E). In addition, the results at 5 and 8 weeks old also showed that the treated mice had a clear preference for the safe zone in both tests. As expected, there was a decrease in preference for the safe zone at 8 weeks compared with the treated mice at 5 weeks, corresponding to the OCT results (Fig. S5B and S5C). These data suggest that *in vivo* base editing could improve vision in *rd10* mice.

**Fig. 5.**
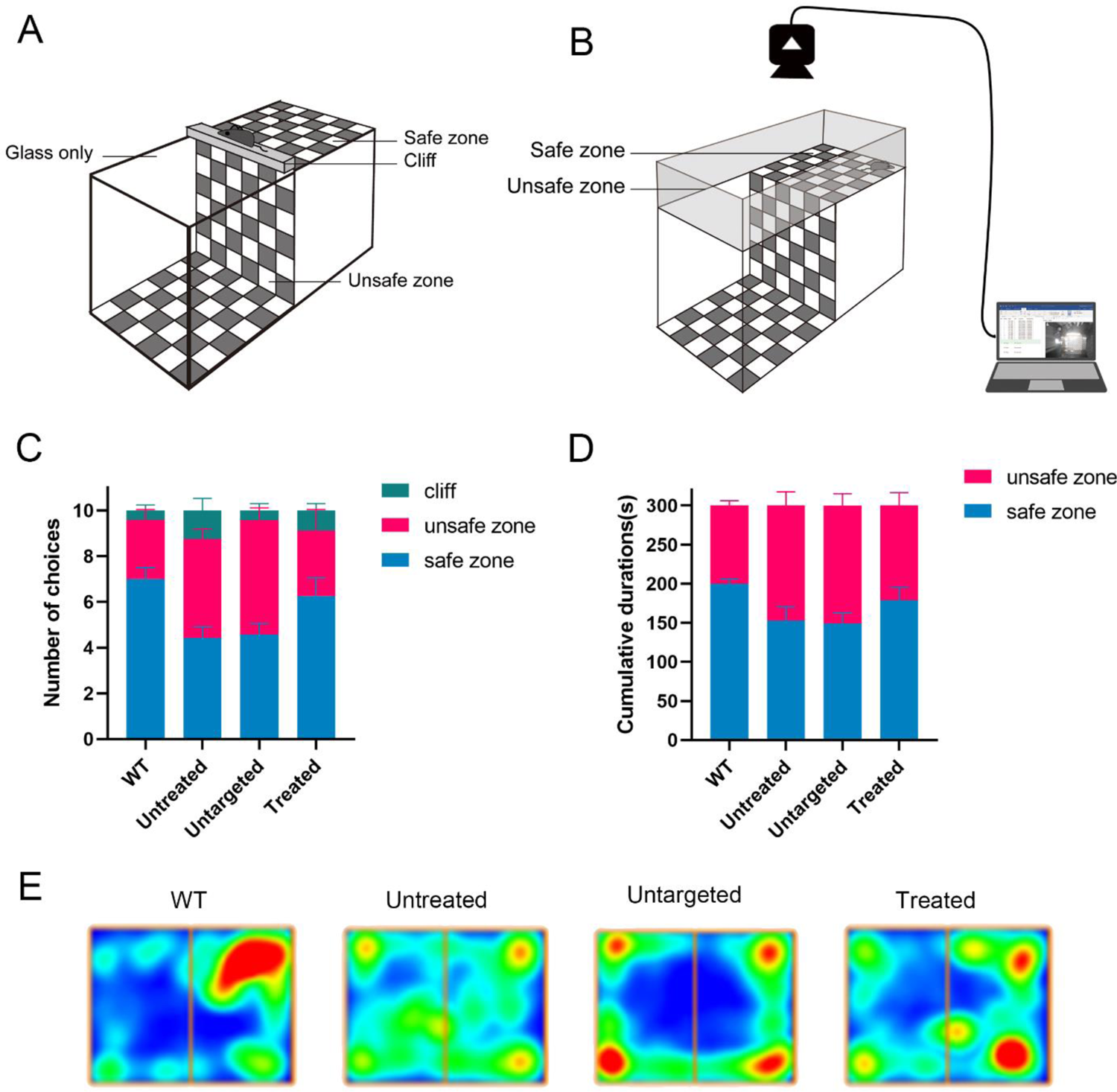
Improvement of visual behavior in treated *rd10* mice by base editing. (**A, B**) Schematic diagrams of two visual cliff tests, A is the cliff step-down test and B is the cliff open field test. (**C**) Quantitative analysis of 10 step-down choices in 12-week-old mice. (**D**) Analysis of the time spent by 12-week-old mice in the safe and unsafe zone. (C, D) Data are shown as mean ± SEM, n=10 for each group. (**E**) The representative heat map of mice staying in the safe zone (right) and unsafe zone (left) in the cliff open field test was analyzed by ANY maze software.

### *In vivo* base editing restores the expression of phototransduction-related genes in *rd10* mice

To evaluate the effect of base editing on the photoreceptor transcriptome, we performed high throughput RNA-sequencing (RNA-seq) on the retinas of 12-week-old WT mice, treated and untreated *rd10* mice (n=3 retinas per group). The correlation analysis between replicates showed that the correlation between samples within each group was high, while the correlation between groups was low (Fig. S8A). We analyzed the differential genes between untreated *rd10* and WT mice (differential expression (DE) analysis: |log2(fold change) | > 1 and p < 0.05) and found that the expression of most photoreceptor-specific genes in untreated mice decreased significantly, which was consistent with a previous report(*27*) (Fig. 6A). The DE analysis revealed 1091 differentially expressed genes in the treated group compared with the untreated group, of which 520 were up-regulated and 571 were down-regulated (Fig. 6B). Gene Ontology (GO) analysis showed that most of these differential genes were enriched in the cascade reactions related to phototransduction (Fig. S8B). Among the up-regulated genes in the treatment group, we observed a remarkable rescue of *Pde6b* gene expression compared to the untreated group (p =1.86×10^-63^). In addition, the expression of rod-specific genes (*Cnga1* and *Rho*), cone-specific genes (*Arr3*, *Opn1mw* and *Pdc*) and key transcription factors related to phototransduction (*Nrl*, *Crx* and *Nr2e3*) was also significantly up-regulated (Fig. 6C). Notably, 18 of the top 20 genes with the highest levels of upregulation were genes involved in phototransduction (Fig. 6D). Simultaneously, we observed that treated mice down-regulated a series of genes associated with biological processes related to photoreceptor cell death, which were significantly enriched in untreated mice, including the calcium signaling pathway (*Prkcb*), cGMP-PKG signaling pathway (*Adcy5*), and neuroactive ligand-receptor interaction (*Lrrk2*, *Gria2*) et etc. (Fig. S8C). These data suggest that base editing can rescue the photoreceptor transcriptome, which is important for both photoreceptor cell survival and phototransduction cascades.

**Fig. 6.**
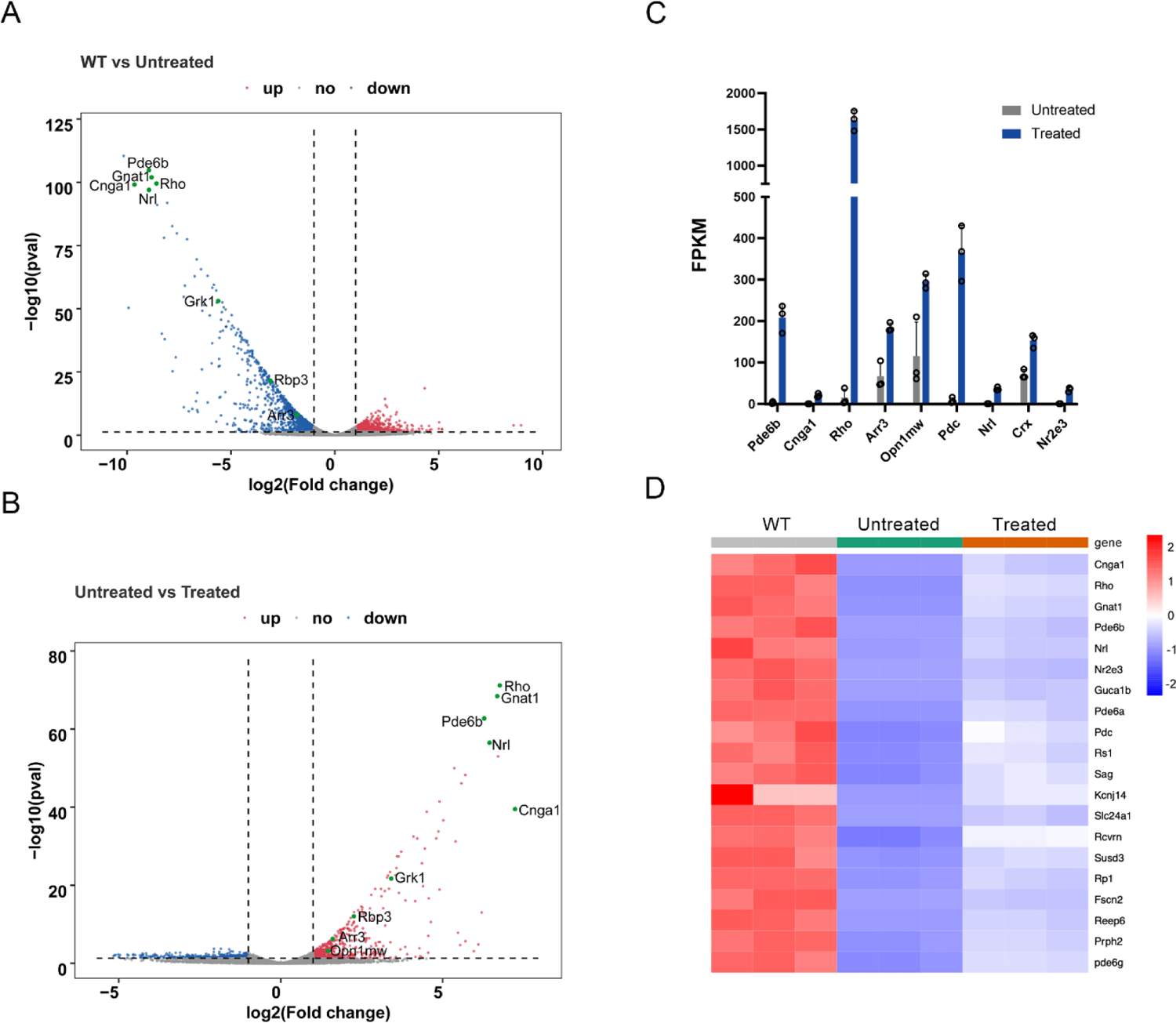
RNA-seq shows the restoration of phototransduction-related gene expression in treated *rd10* mice. (**A**) Volcanic plot of differential expression between WT mice and untreated *rd10* mice (|log2(fold change) |)> 1 and p < 0.05; red: up-regulated; blue, down-regulated). (**B**) Volcanic plot of differential expression between untreated mice and treated *rd10* mice. (**C**) The expression of key genes related to photoreceptors and phototransduction was significantly upregulated. Mean ± SD are shown, n=3 for each group. (**D**)The heat map of the top 20 genes that were most significantly upregulated in the treated group compared to the untreated group, n=3 for each group.

## Discussion

In recent years, base editing has provided a promising treatment for some rare monogenic diseases with no or limited treatment. Inherited retinal diseases (IRDs) are well suited for the application of this approach because most of them are caused by monogenic mutations, such as LCA, Stargardt disease and RP(*28*). Herein, we demonstrated that AAV8-mediated split-intein ABE could effectively correct the pathogenic mutation of photoreceptors and protect against photoreceptor degeneration in the *rd10* mouse model of RP. Our study provides a framework for the preclinical development of a base-editing therapeutic for other RPs caused by different mutations.

Photoreceptor degeneration occurs rapidly and dramatically in RP, thus early and effective treatment is essential to prevent the development of serious manifestations. In the past, most therapeutic strategies for *rd10* mice have focused on the mechanisms of photoreceptor death, such as inhibition or activation of related pathways, neuroprotective treatment and antioxidant treatment(*29–32*). These strategies have certain protection against photoreceptors but only maintained short-term therapeutic effects without genetic correction. Furthermore, only the cone cell bodies remained in the late stages of the disease in these studies, while the rod was lost. In this study, we used AAV-mediated base editor to irreversibly correct pathogenic gene. Notably, the correction efficiency of the target site in retina was up to 37.41% (average 29.63%) at the DNA level and was up to 91.95% (average 89.79%) at the cDNA level, which effectively preserved rod and cone photoreceptors (Fig. 2). In *Pde6b*-RP, the death of mutant rods leads secondarily to the death of cones and, subsequently, severe vision loss; consequently, rod rescue is critical for long-term treatment. The length of rod OS in the treated group was 33% of that in WT mice compared with untreated mice and the scotopic ERG showed considerable a- and b-wave amplitude recovery, indicating a partial restoration of rod-driven retinal (Fig. 4A). Studies have shown that rod rescue has a protective effect on the cone survival and function(*33, 34*). Our data also found that approximately 89% of cones were retained in treated mice at 12 weeks of age, and both M-cone and S-cone mediated retina function were preserved (Fig. 2C, 4B and 4C). In addition, base editing effectively prevented the degeneration of the retinal structure and function. However, in non-invasive OCT and behavioral follow-up results, we found that the retinal structure and function of the treated mice decreased significantly at 8 weeks compared to 5 weeks, and only slightly decreased at 12 weeks (Fig. S6). We reasoned that this is because photoreceptors have died or are dying before the pathogenic mutation can be effectively corrected and dying cells cannot be rescued. To rapidly and effectively prevent photoreceptor degeneration in *rd10* mice, Paola Vagni et al. performed gene editing therapy through *in vivo* electroporation at P1 and P3(*13*). Although pre-onset treatment of *rd10* rescued photoreceptor degeneration, injection prior to P14, especially P7, was highly traumatic to the eyes of mice.

In recent decades, gene augmentation therapy for *Rpe65*-LCA has achieved encouraging success in animal models and the clinic, but several clinical studies have also shown that the gene therapy effect does not last “forever”(*35–37*). By contrast, AAV-mediated base editors can irreversibly correct the pathogenic genes and might have a sustained therapeutic effect. However, the bystander effects of base editing in clinical application remains a concern. Currently, several strategies have been developed to reduce the bystander effect, such as narrowing the editing window of the base editors(*38*), limiting the expression time of the base editors *in vivo*(*39*), and restricting the connection between deaminase and Cas9(*40*). We believe that the bystander effect problem will be solved in the future. In this study, as in the previously reported a *Rpe65-rd12* base editing study(*18*), the positive effects of high levels of editing at the target site were much higher than the possible negative effects of the bystander despite the bystander editing.

In conclusion, we demonstrate that AAV-mediated base editor delivery can effectively correct pathogenic mutation and protect against photoreceptor degeneration and visual function impairment in *rd10* mice. As in this study, base editing provides a treatment method for incurable RPs and other IRDs caused by mutations affecting photoreceptors. This work expands the disease spectrum that base editors can treat and paves the way for base editor clinical translation.

## Materials and Methods

### Cell line generation

The pX330 plasmid was used to construct CRISPR/Cas9 vectors for generating the HEK293-*Pde6b* mutant cell line. HEK293 cells were transfected with CRISPR/Cas9 plasmid and a donor plasmid containing the PDE6B-R560C-PURO sequence. The transfected cells were screened for puromycin, and the monoclonal cells were extracted to 96-well plates for amplification and culture. Then the genomic DNA of the cells was collected for PCR and sequencing verification. Finally, the HEK293-*Pde6b* mutant cell line was generated by stably integrating the PDE6B-R560C sequence into the AAVS1 locus. The primers constructed by mutant cell lines are listed in Table S1.

### Plasmid Construction

VRQR-ABEmax (#119811), xCas9(3.7)-ABE(7.10) (#108382), xABEmax (#119813), NG-ABEmax (#124163), SpG-ABE8e (#140002), NG-ABE8e (#138491) and pSPgRNA (#47108) plasmids were purchased from Addgene (Watertown, MA). To generate the ABE8e-SpG plasmid, NG-ABE8e was digested by *NotI* and *EcoRV* and subcloned into the SpG-ABE8e plasmid backbone by In-Fusion cloning (Takara Bio, Mountain View, CA). SgRNA-A6 and sgRNA-A8 targeting the A→G R560C mutation site on exon 13 of *Pde6b* gene in the mouse genome were designed by an online webtool (https://benchling.com). All sgRNAs constructed were generated by T4 ligation of annealed oligos into the *BbsI* digested pSPgRNA plasmid. Both the N-NG-ABE8e and C-NG-ABE8e.sgRNA-A8 vectors used the CBh promoter and were generated by In-Fusion cloning of PCR amplification inserted into restriction enzyme-digested backbones. All constructed plasmids were verified by sequencing.

### Cell culture and transfection

HEK293-*Pde6b* mutant cell lines were maintained in DMEM medium supplemented with 10% fetal bovine serum (FBS) and cultured at 37℃ with 5% CO2. The HEK293-*Pde6b* mutant cells were seeded into 24-well plates at 1×10^5^ cells per well 24 hours before transfection. Six adenine base editors (VRQR-ABEmax, xCas9(3.7)-ABE(7.10), xABEmax, NG-ABEmax, SpG-ABE, NG-ABE8e and ABE8e-SpG) were co-transfected with sgRNA-A6 or sgRNA-A8 into HEK293-*Pde6b* mutant cell lines, respectively. ABE plasmids and sgRNA plasmids were transfected at 750ng and 250ng, respectively. Cells were harvested for genomic DNA extraction 72 hours after transfection.

### AAV Vector Production

AAV8.N-ABE and AAV8.C-ABE.sgRNA-A8 were obtained by packaging N-NG-ABE8e and C-NG-ABE8e.sgRNA-A8 into AAV8 vectors. All AAV8 vectors were produced by triple plasmid transfection of HEK293 cells (ATCC, Manassas, VA) as previously described(*41*). The genome titer (GC/ml) of the AAV8 vector was determined by digital droplet polymerase chain reaction (ddPCR) using forward primer 5’-TAGTTGCCAGCCATCTGTTG-3’, reverse primer 5’-TAGGAAAGGACAGTGGGAGT-3’, and probe 5’-Fam-CCCGTGCCTTCCTTGACCCT-BHQ-3’. All vectors used in this study passed the endotoxin assay using the end-point chromogenic endotoxin test kit (Xiamen Bioendo Technology Co.,Ltd., Xiamen, China).

### Animals

*Rd10* mice (B6.CXB1-Pde6bRd10, Stock No: 004297) were purchased from Jackson Laboratory (Bar Harbor, Maine). The background of the wild-type mice used in this study was C57BL/6J. *Rd10* mice in late pregnancy were moved to a dark room in total darkness. Newborn *rd10* mice were moved out of the dark room when they were 4 weeks old and then raised on a 12h-12h light-dark cycle. Unless otherwise stated, mice were anesthetized with intraperitoneal ketamine (80 mg/kg) and xylazine (12 mg/kg) in this study. The pupils were dilated with an eye drop containing 0.5% tropicamide and 0.5% phenylephrine hydrochloride. All animal protocols were approved by the Institutional Animal Care and Concern Committee at Sichuan University, and animal care was in accordance with the committee’s guidelines.

### Subretinal injection

*Rd10* mice were bilaterally and subretinally injected at 2 weeks of age. After inhalation of isoflurane and pupil dilation anesthesia, a limbal incision was made under a stereo microscope with a 31-gauge needle. Then, a 33-gauge blunt needle (Hamilton, Reno, New York) was inserted through the incision and directed into the subretinal space, avoiding damage to the lens. Each eye received a mixture of AAV9.N-ABE and AAV8.C-ABE.sgRNA-A8 or AAV8.C-ABE.sgRNA-ctrl at 1:1 (3×10 ^9^ GC /eye for each vector) in a volume of 1 μl, as described. Immediately after injection, a retinal imaging microscope (Micron IV, Phoenix Research Labs, Pleasanton, California) was used to observe the fundus and evaluate whether the injection was successful.

### Electroretinography (ERG)

Scotopic ERG and photopic ERG were measured at 5 w and 12 w in mice. ERG was recorded according to the manufacturer’s instructions for the Phoenix Ganzfeld ERG (Phoenix Research Laboratories, Pleasanton, CA). All animals underwent dark adaptation overnight before ERG recording. After anesthesia, the mice were placed on the heating pad, and ERG was recorded after pupil dilation. The reference electrode was placed subcutaneously in the forehead between the ears, and the ground electrode was placed subcutaneously in the tail. Corneal electrodes were placed on the cornea after application of 2.5% Hypromellose. Scotopic ERG was recorded at the following increasing stimulus intensities of −1.1, 0.1, 0.7, 1.0, 1.3 and 2.2 log cd s/m^2^. Photopic ERGs were stimulated with green and UV light, and photopic recordings were performed after a 5-min light-adaptation interval at a background light intensity of 1.3 log cd s/m^2^, which also served as background light for the duration of the optical recording. Photopic ERG was recorded at the following increasing stimulus intensities of 0.3, 0.9, 1.5, 2.1, 3 and 3.9 log cd s/m^2^.

### Optical coherence tomography (OCT)

OCT images were acquired with Spectralis HRA+OCT (Heidelberg engineering, Heidelberg, Germany) and a 30-angle lens was used to determine in vivo retinal thickness. For quantification, 360-degree scans were performed at 15-degree intervals from the vertical direction with the optic disc as the center. Retinal and ONL thicknesses were measured at 6 inferior and 6 superior positions (0.25, 0.5.1.0, 1.5, 2.0, 3.0 mm from the optic disc) using OCT system software, and the thickness of each position was averaged for 45 degrees clockwise and counterclockwise, vertical and horizontal directions.

### Western blot analysis

Western blot analyses were performed on retinal lysates from mouse eye. The mouse eye retina was dissected and placed in a microcentrifuge tube containing 50 μl RIPA buffer. The total protein concentration was determined by BCA protein assay (Thermo Scientific; Waltham, MA). PDE6B protein was detected by Rabbit anti-PDE6B antibody (1:500; Thermo Scientific, PA1-722). Mouse anti-GAPDH antibody (1:10000; ABclonal; Cat# AC002) was used to detect GAPDH. Blots were imaged and analyzed by iBrightTM CL1000 imaging systems (Thermo Scientific; Waltham, MA).

### Immunohistochemistry

Eyes were enucleated and fixed in 4% paraformaldehyde/PBS at room temperature (RT) for 1 h. The eyeballs were dehydrated in 30% sucrose solution for 2 h and embedded with O.C.T. Cryosections (10 μm) were cut in sagittal orientation, rinsed with PBS for 5 min, and blocked with 5% normal goat serum and 0.2% TritonX-100 in PBS for 1 h at RT. Then, the sections were incubated with primary antibodies in blocking solution overnight at 4 °C. The primary antibodies used in this study were as follows: rabbit anti-PDE6B (1:500; Thermo Scientific; PA1-722), mouse anti-rhodopsin (1:1000; Sigma Aldrich; Clone RET-P1), and rabbit anti-cone arrestin (1:1000; Millipore; AB15282). After washing with PBS 3 times, sections were incubated with corresponding secondary antibodies for 1 h at RT, including goat anti-rabbit Alexa Fluor 594 (1:1000; Abcam; ab150080), goat anti-rabbit Alexa Fluor 488 (1:1000; Abcam; ab150070) and goat anti-mouse Alexa Fluor 594(1:1000; Abcam; ab150116). Sections were stained with DAPI and imaged with a confocal laser microscope (Nikon, Tokyo, Japan).

### Immunostaining of RPE and Retina Flatmounts

For RPE and retinal flatmounts, after enucleation, eyes were marked on the 12 o’clock point (dorsal) of the limbus and the cornea, lens, vitreous and optic nerve bud were carefully removed to generate eye cups. Then the whole retina and RPE/choroid/sclera were carefully separated, and four radial cuts were made toward the optic nerve head to flatten the RPE/choroid/sclera or retina. The RPE/choroid/sclera or retina was fixed in 4% paraformaldehyde in 0.1 M PBS for 1 h at RT and then washed in PBS 3 times (10 min each). Then, the specimens were incubated in 10% normal goat serum (NGS) blocking solution for 1 hour at RT, and staind with primary antibody for 48 h at 4 °C. The primary antibodies used for RPE/choroid/sclera was rat anti-ZO-1 (1:100; Santa Cruz; sc-33725); those used for retina were rabbit anti-S opsin (1:50; Novus; NBP1-20194) and rabbit anti-M opsin (1:1000; Novus; NB110-74730). Then, the specimens were washed in PBST 3 times (15 min each) and incubated with secondary antibodies for 1 h at RT. The secondary antibodies used in this study were as follows: goat anti-rabbit IgG Alexa Fluor 594 (1:1000; Abcam; ab150080) and goat anti-rat IgG Alexa Fluor 488 (1:1000; Abcam; ab2534074). The specimens were then washed in PBS 3 times (15 min each) and imaged with a confocal laser microscope (Nikon, Tokyo, Japan). The RPE flatmounts were stained with DAPI before photography.

### Transmission electron microscopy (TEM)

Eyes were enucleated and prefixed with a 3% glutaraldehyde. Then, the tissue was postfixed in 1% osmium tetroxide, dehydrated in series acetone, infiltrated in Epox 812 for a longer and embeded. The semithin sections were stained with methylene blue and ultrathin sections were cut with diamond knife, stained with uranyl acetate and lead citrate. Sections were examined with JEM-1400-FLASH Transmission Electron Microscope (JEOL, Tokyo, Japan).

### Base editing analysis with Sanger sequencing and EditR software

On-target genomic regions of interest were amplified by PCR and analyzed with Sanger sequencing. Then, the sequencing graphs were further quantified by EditR software (baseditr.com)(*42*). The primers used for amplifying target loci are listed in Table S1.

### Amplification and NGS of genomic DNA and mRNA samples

To evaluate the editing effect of DNA and mRNA in mouse eyes, genomic DNA and total RNA of retinal tissue were extracted with QIAamp DNA Micro Kit (Qiagen, 56304) and RNeasy Plus Micro kit (Qiagen, 74034), respectively. Then, RNA was reverse transcribed to cDNA using the PrimeScript™ RT reagent Kit with gDNA eraser (Takara, Kusatsu, Japan) according to the manufacturer’s instructions. Subsequent nested PCRs were performed to generate amplicons for NGS and the second PCR product was purified using GeneJET Gel Extraction Kit (Thermo Fisher Scientific). Furthermore, the top eight potential off-target sites for sgRNA8 were identified by the algorithm described in www.benchling.com (Table S2). These off-target sites were amplified by nested PCR in retina tissue genomic DNA and deep sequenced with NGS. The primers used for amplifying each target loci are listed in Table S3. Libraries were made from the second PCR products and sequenced on Illumina NovaSeq (2 × 250 bp paired end, Personal Biotechnology Co., Ltd, Shanghai, China). Data were processed according to standard Illumina sequencing analysis procedures. Processed reads were mapped to the expected PCR amplicons as reference sequences using custom scripts. Reads that did not map to reference were discarded. Indels were determined by comparison of reads against references using custom scripts.

### H&E staining

Eyes were enucleated and fixed in 4% paraformaldehyde at RT for 6 h, dehydrated through an ethanol series and xylene, and then embedded in paraffin. H&E staining was performed on 5 µm sections from paraffin-embedded retinas according to standard protocols for light microscopy and photomicrography (Leica, Wetzlar, Germany).

### Behavioral test

The two visual cliff tests were performed on the visual acuity of mice under 10 lux light intensity. The experimental equipment consists of a platform with a plaid pattern and a chamber with a transparent glass bottom on it. The chamber is divided by a plaid pattern into a deep plaid area and a transparent light area. For the step-down trials, a rising platform was placed between the grid area and the transparent area to create the illusion of a cliff. Each mouse was placed on the middle ascending platform, allowing them to walk to the shallow side (safe choice) or the deep side (unsafe choice). Each mouse performed 10 trials, and if the mouse moved towards the lattice area, the choice was safe; if the mouse moved towards the clear area, the choice was not safe, and the mouse’s choice was recorded each time. For the cliff open field test, the raised platform was removed, the mouse was placed on the platform, and record the time it spent in the safe and unsafe zones, as well as the number of shuttles. Collection and analysis of open field test data were performed using ANY maze software.

### Statistics

GraphPad Prism9 was used to perform all data analyses. Data are presented as mean ±SD in Fig.1B, 1E, 2C, 2E, 6 and S2A. Data are presented as mean ± SEM in Fig.1G, 3B, 4, 5, S5 and S7. One - way ANOVA with Tukey’s post-hoc test was used in Fig. 2B and 2C. Two-way ANOVA with Tukey’s post-hoc test in Fig. 4A-4C. In all tests, p<0.05 was considered significant.

## Acknowledgements

We thank Chengdu Genevector Therapeutics Inc. (Cheng Du, China) for providing professional technical support.

## Author contributions

Y.Y. and F.L conceived this study and designed the experiments; R.L. constructed the plasmid vectors; X.J. produced the AAV8 vector and endotoxin assays; J.S. and L.S. performed the mouse studies; H.C. performed on the NGS analyses; K.S., Q.Z. and J.X. performed the behavioral tests; J.S. and K.S. performed histopathology assays; J.S. wrote the manuscript; Y.Y., D.C. and Y.W. edited the manuscript. All authors read and approved the final manuscript.

## Data and materials availability

All data needed to evaluate the conclusions in the paper are present in the paper and/or the Supplementary Materials. Additional data related to this paper may be requested from the lead contact Y.Y.

## Supplementary Materials

**Fig. S1.**
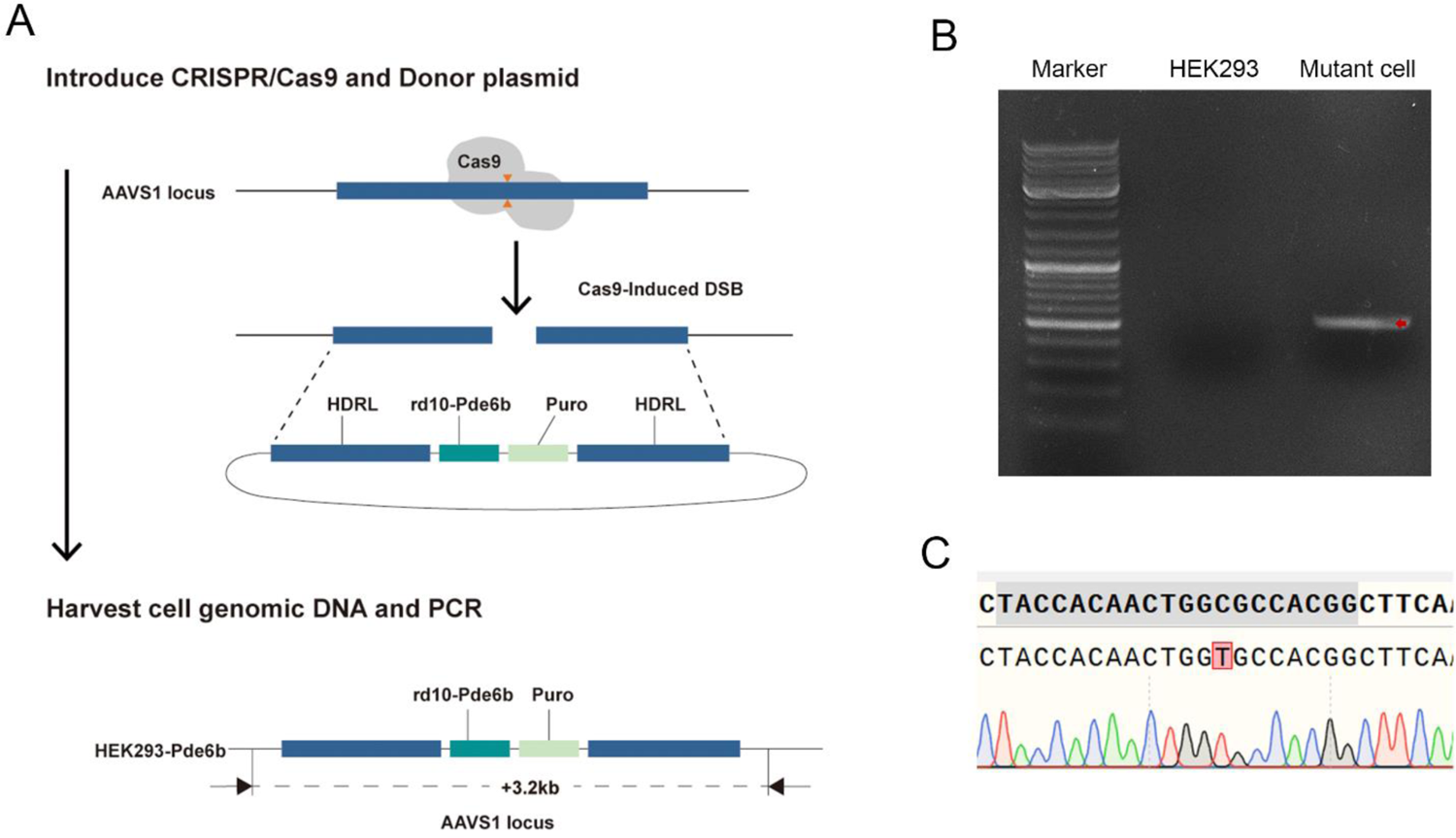
Construction of HEK293-*Pde6b* mutant cell line using CRISPR/Cas9. (A) Schematic diagram of HEK293-*Pde6b* mutant cell line construction. (B) Gel electrophoresis verified that the mutant sequence was successfully inserted into the DNA genome of HEK293 cells. The inserted mutant sequences are marked in red. (C) The successful construction of HEK293-*Pde6b* mutant cell line was verified by Sanger sequencing. The red square is the mutation site.

**Fig. S2.**
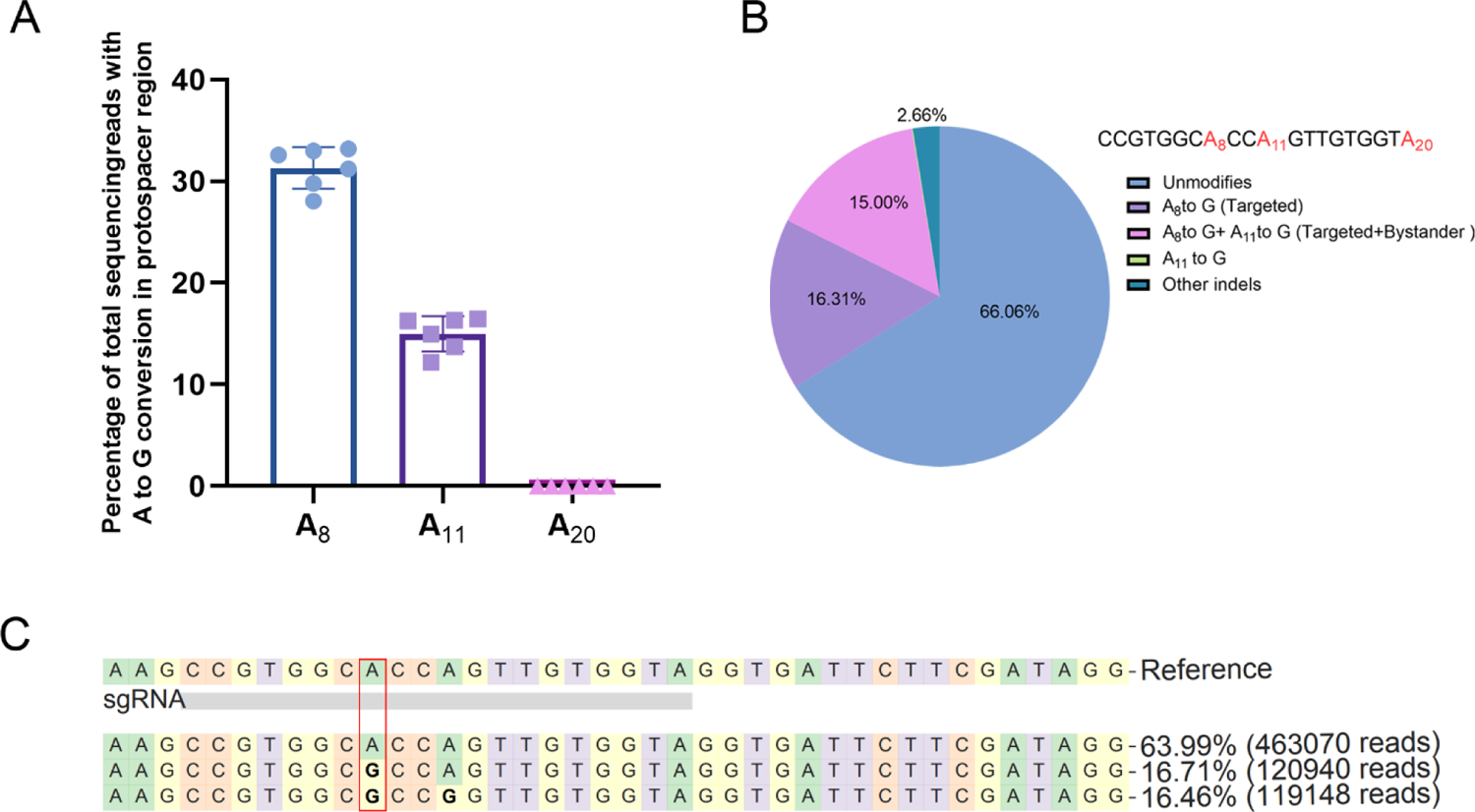
*Pde6b* base editing analysis in high dose treatment group. (A) Editing efficiencies at different A positions within the protospacer region were determined by NGS. A8 is the target site. Data are shown as mean ± SD, n=6. (B) Pie chart shows the average composition of allele variants at DNA level in treated *rd10* mice. (C) Representative sequence alignment showing top 2 sequences of base editing outcomes in *Pde6b* locus after high-dose treatment. The red box marks targeted site.

**Fig. S3.**
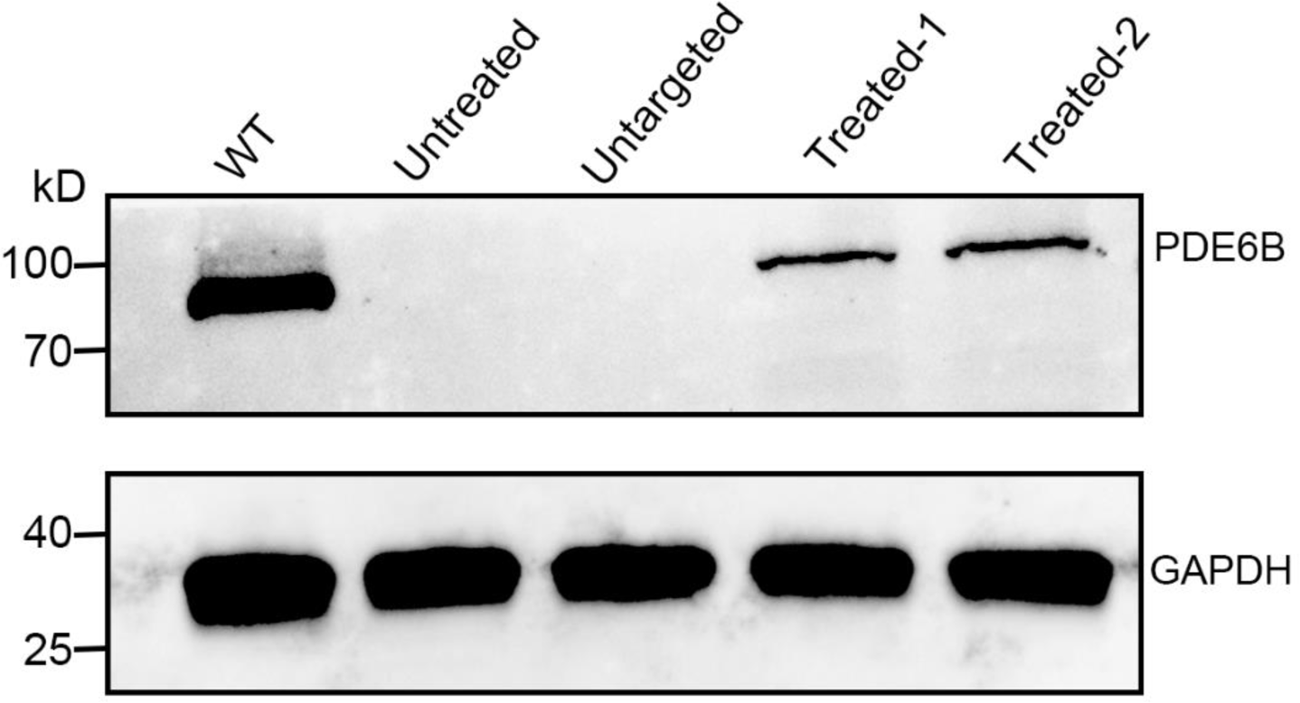
*Pde6b* gene expression by Western blot analysis. The expression of PDE6B protein was detected in the retinal lysate of mice at 12 weeks of age. PDE6B protein (98 kDa); GAPDH protein (36 kDa).

**Fig. S4.**
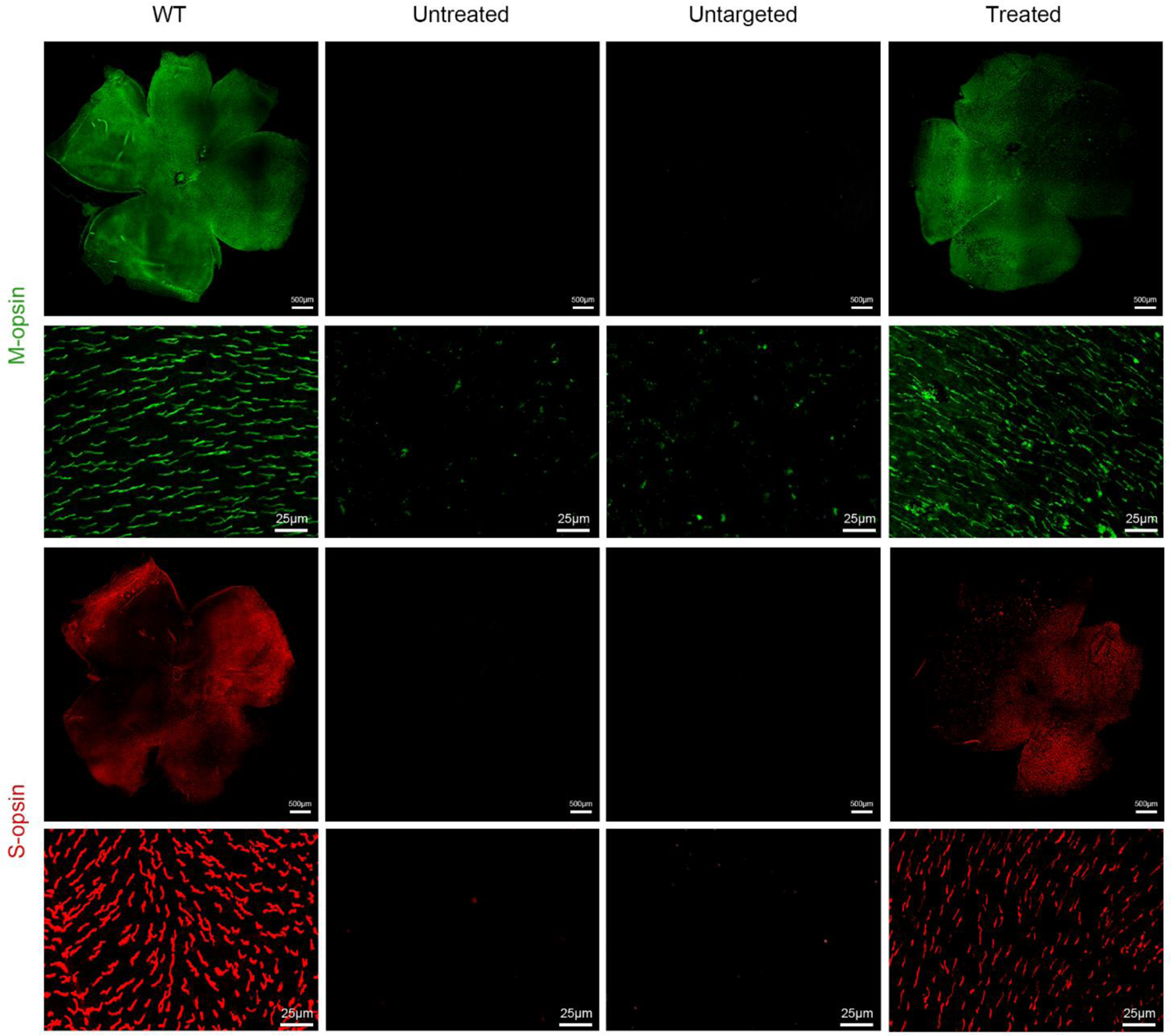
Representative images of retinal flatmounts immunolabeled with specific M-cone and S-cone antibody. M-cone, green; S-cone, red. Overall view of the retinal flat-mounts in first and third row. Magnified view of M-cone and S-cone in second and fourth rows. The S-cone and M-cone were clearly observed in the treated mice, while S-cone and M-cone were lost in untreated and untargeted mice. Scale bars: 500 μm for low power; 25 μm for high power.

**Fig. S5.**
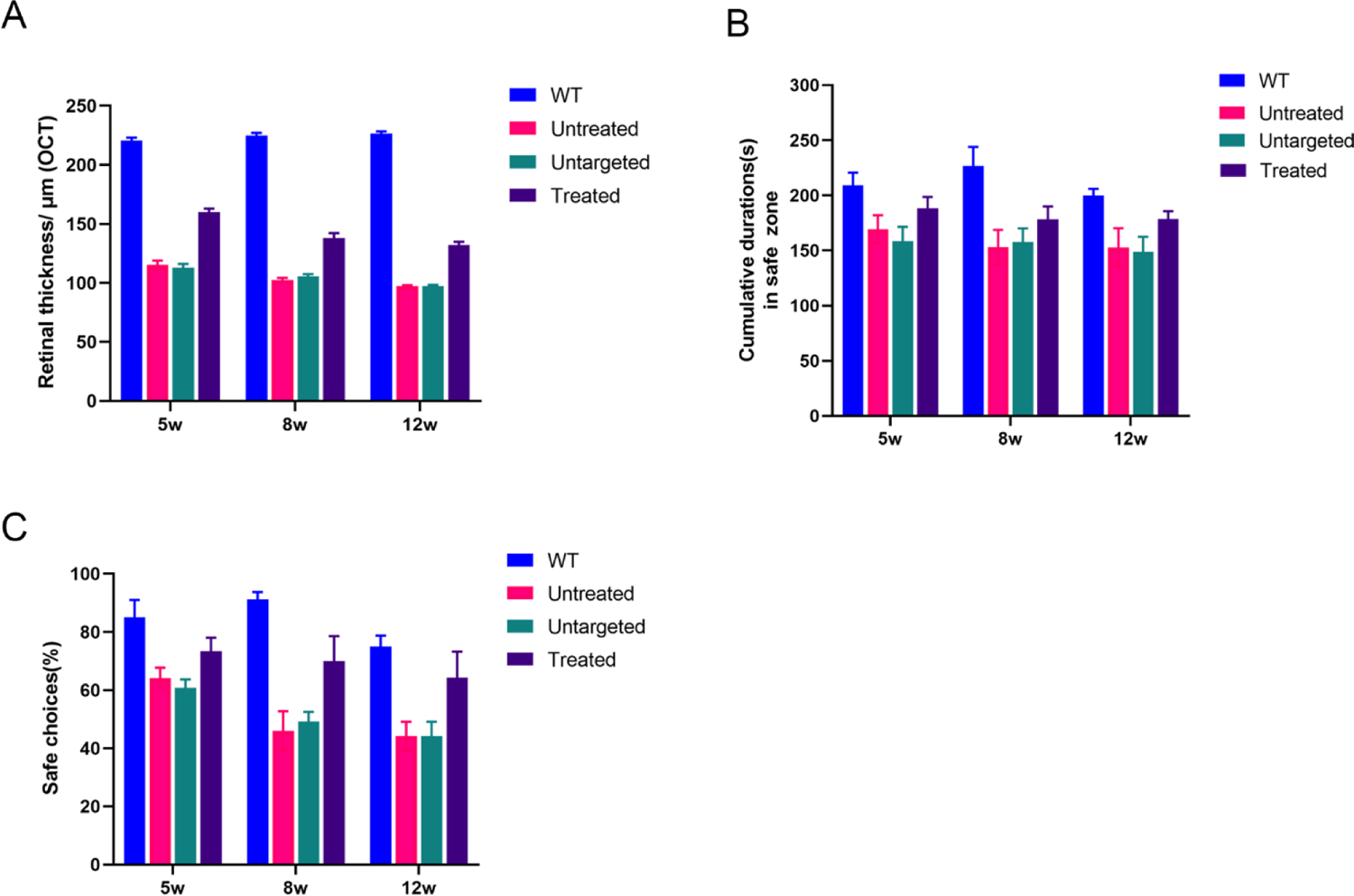
Time course of base editing treatment efficiency at 5, 8 and 12 weeks. (A) Quantitative results of retinal thickness at different time points obtained by OCT (n=10, each group). Mean ± SEM are shown. (B) The results of mice’s preference for safe zone at different time points detected by cliff step-down test. (C) Analysis of the time spent by mice in the safe zone at different time points detected by the cliff open field test. (B, C) Data are shown as mean ± SEM, n=10 per each group.

**Fig. S6.**
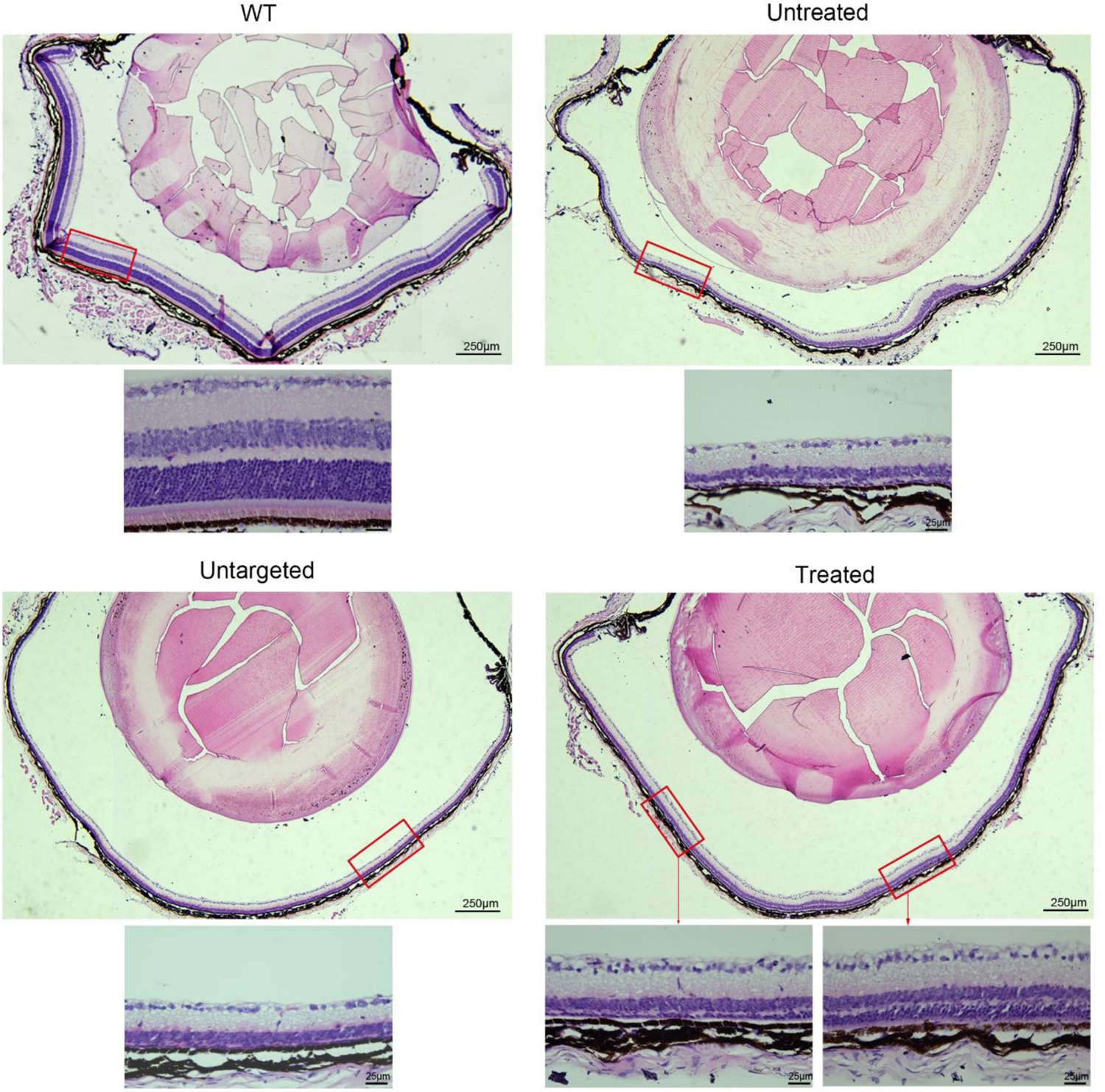
Representative photomicrographs of retinas in mice at 12 weeks of age. Histological analysis of the eyes at 12 weeks of age by hematoxylin and eosin stain. The ONL in untreated mice disappeared, and the ONL layer of more than 3/4 of retinas was retained in treated mice. Scale bars: 250 μm for low power; 25 μm for high power.

**Fig. S7.**
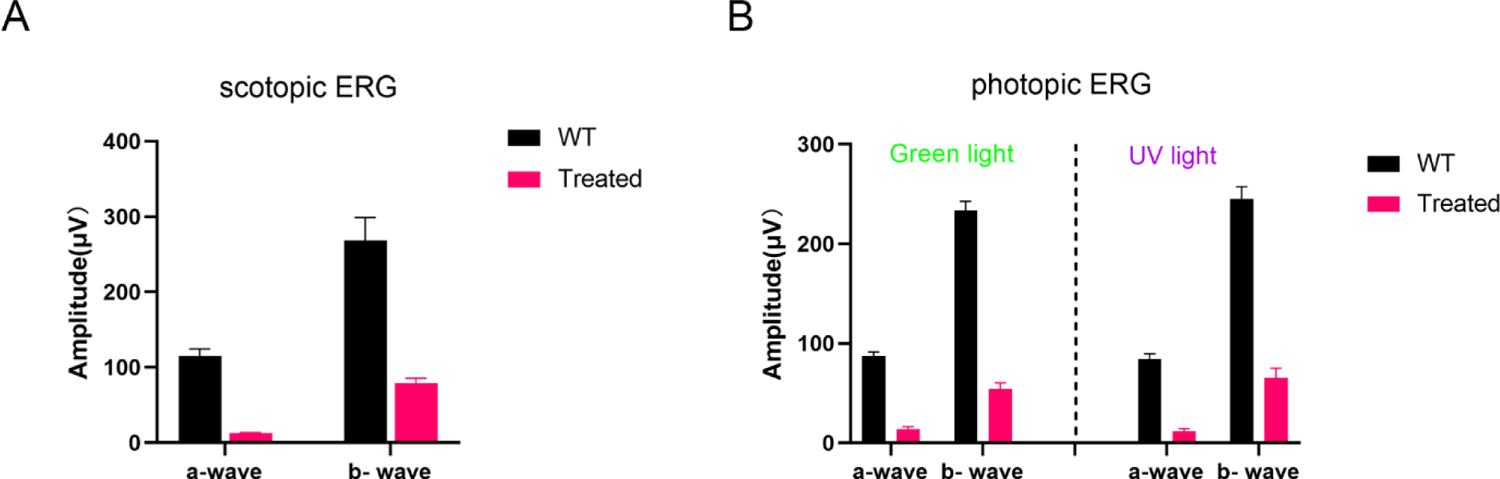
Comparison of scotopic and photopic ERG responses in WT and treated *rd10* mice at 12 weeks of age. (A) Quantification of scotopic ERG a- and b-waves at 2.2 log cd s/m^2^ stimulus intensity in WT and treated *rd10* mice. (B) Quantification of photopic ERG a- and b-waves at the stimulus intensity of 3.9 log cd s/m^2^. left: green light; right: UV light. (A, B) Data are shown as mean ± SEM, n=10 per each group.

**Fig.S8.**
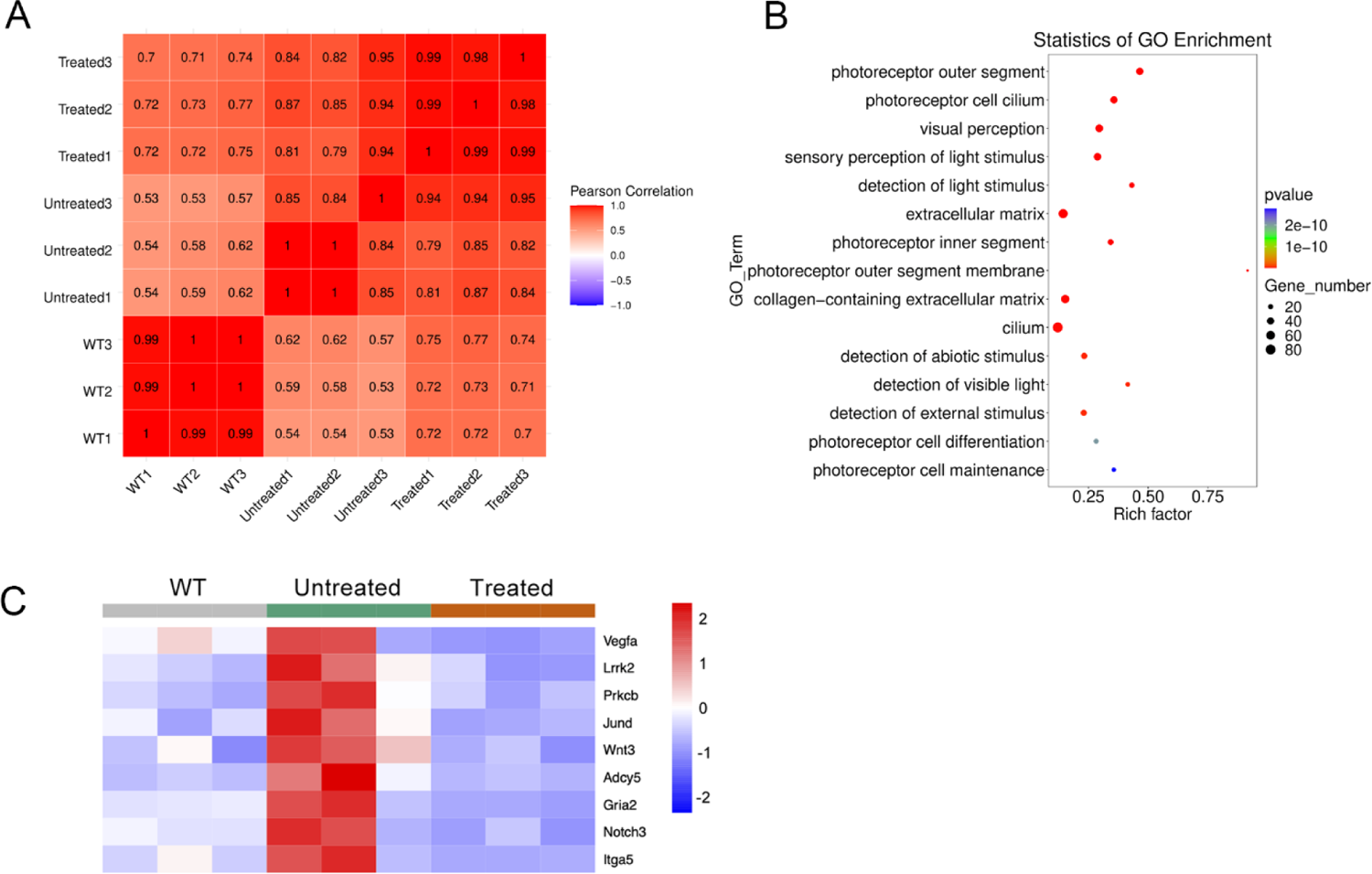
The RNA-seq analysis of retinal from 12-week-old WT, untreated and treated rd10 mice. (A) Heatmap of RNA-seq correlation between each sample by Pearson’s correlation coefficient analysis. (B) GO analysis of differentially expressed genes between the untreated and AAV8-treated group. (C) Heat map shows downregulation of gene expression associated with photoreceptor death cells in the treated group compared to the untreated group.

---

**Table S1.**
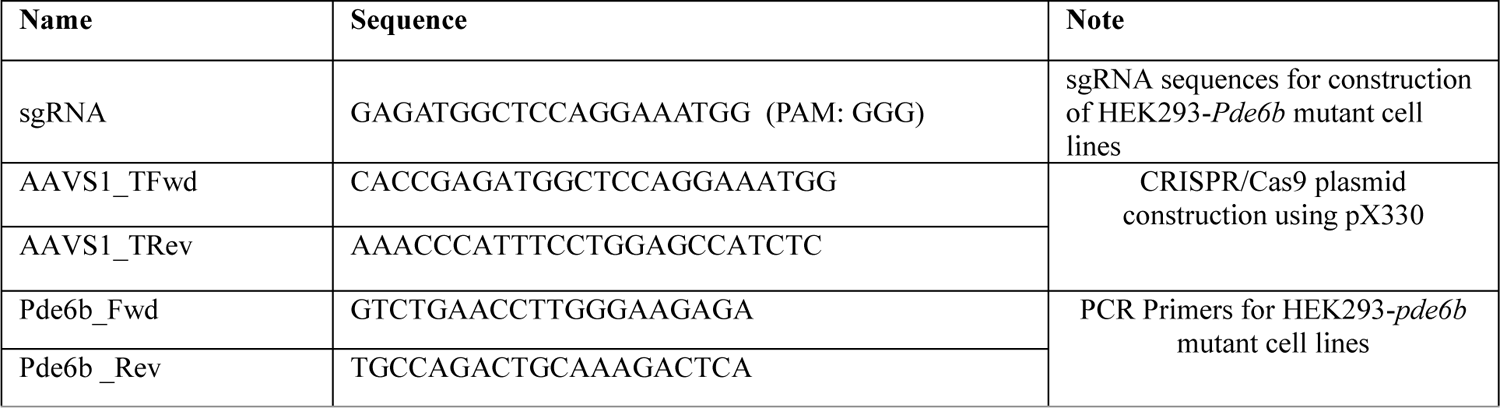
Primers and sequences for construction of HEK293-Pde6b mutant cell lines.

**Table S2.**
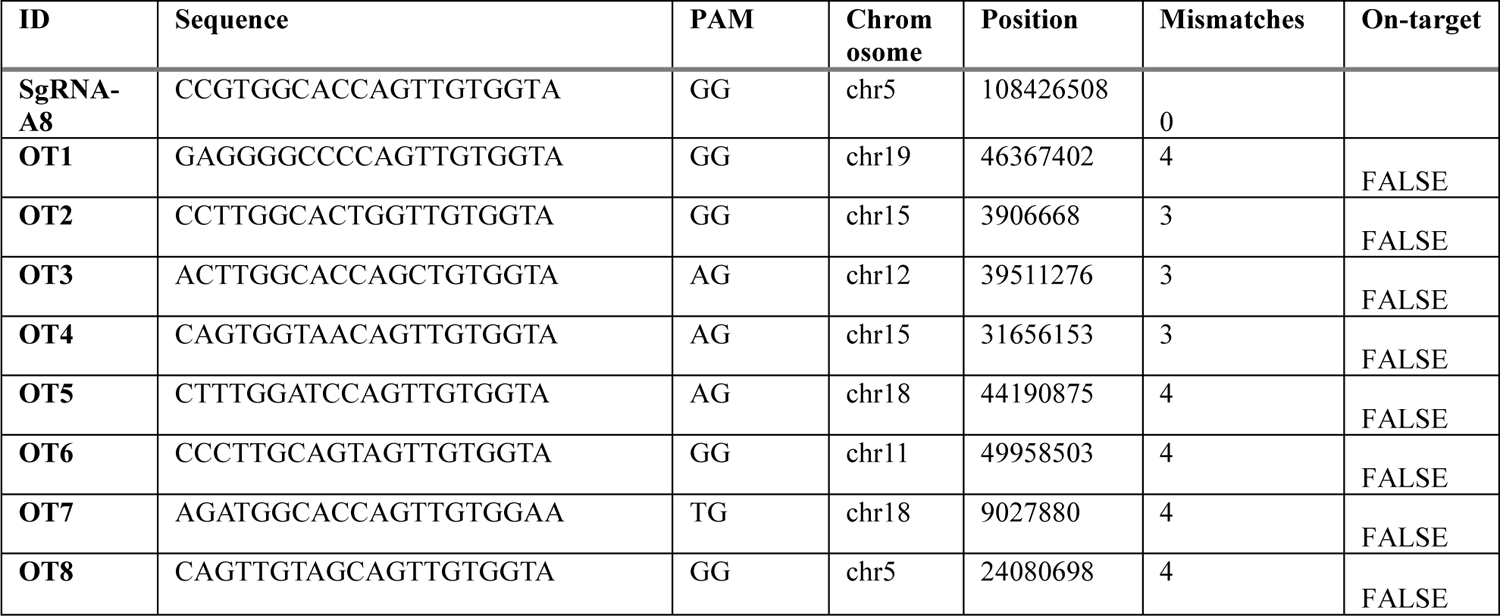
Off-target analysis. Potential off-target sequences for sgRNA-A8 identified and scored by Benchling’s off-target analysis.

**Table S3.**
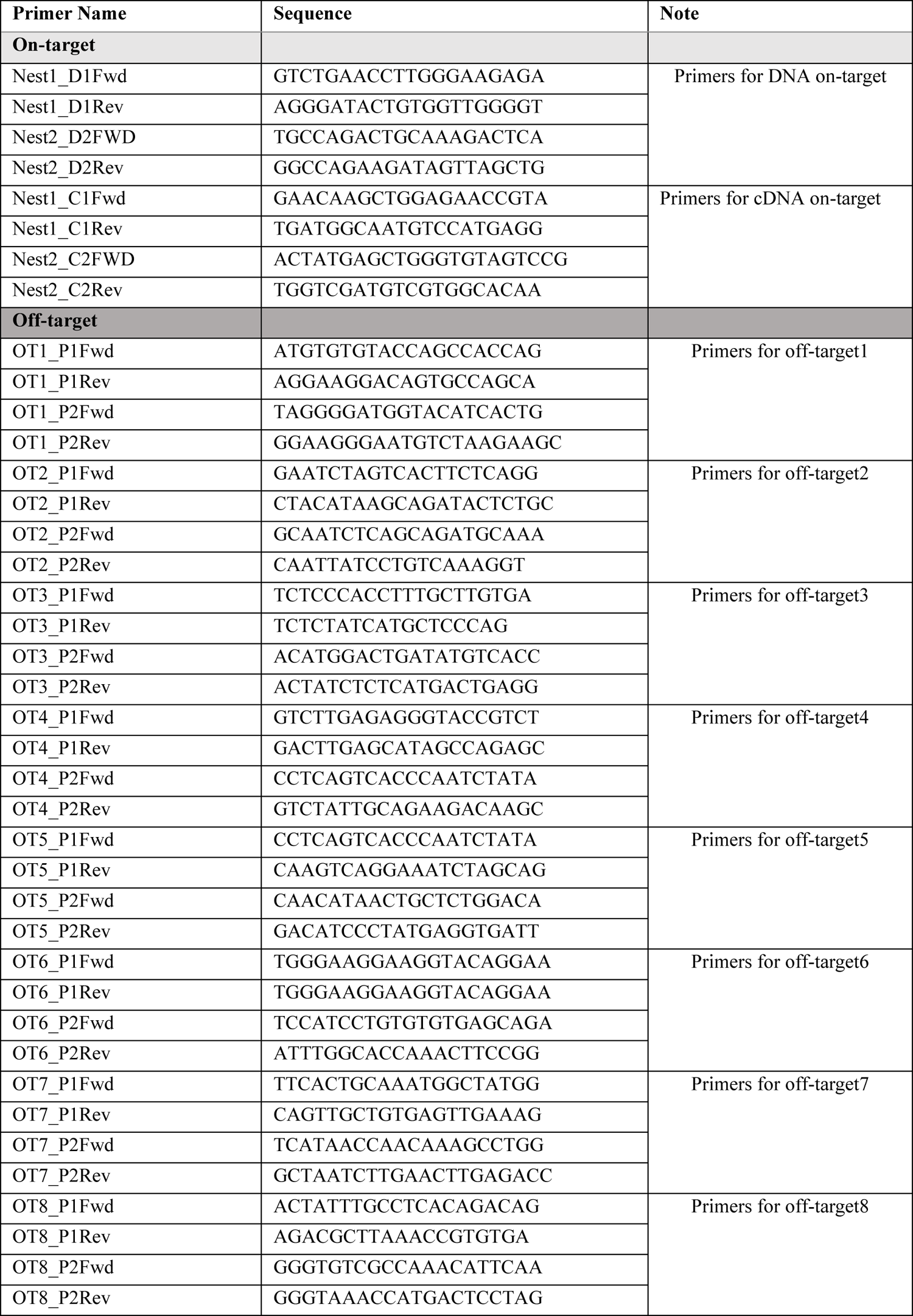
PCR primer sequences for detecting potential on-target and off-target effects by NGS assay.

